# Learning induced neuronal identity switch in the superficial layers of the primary somatosensory cortex

**DOI:** 10.1101/2023.08.30.555603

**Authors:** Jiaman Dai, Qian-Quan Sun

**Affiliations:** Department of Zoology and Physiology, University of Wyoming, Laramie, WY82071, USA.; Wyoming Sensory Biology Center of Biomedical Research Excellence, University of Wyoming, Laramie, WY82071, USA.

## Abstract

During learning, multi-dimensional inputs are integrated within the sensory cortices. However, the strategies by which the sensory cortex employs to achieve learning remains poorly understood. We studied the sensory cortical neuronal coding of trace eyeblink conditioning (TEC) in head-fixed, freely running mice, where whisker deflection was used as a conditioned stimulus (CS) and an air puff to the cornea delivered after an interval was used as unconditioned stimulus (US). After training, mice learned the task with a set of stereotypical behavioral changes, most prominent ones include prolonged closure of eyelids, and increased reverse running between CS and US onset. The local blockade of the primary somatosensory cortex (S1) activities with muscimol abolished the behavior learning suggesting that S1 is required for the TEC. In naive animals, based on the response properties to the CS and US, identities of the small proportion (∼20%) of responsive primary neurons (PNs) were divided into two subtypes: CR (i.e. CS-responsive) and UR neurons (i.e. US-responsive). After animals learned the task, identity of CR and UR neurons changed: while the CR neurons are less responsive to CS, UR neurons gain responsiveness to CS, a new phenomenon we defined as ‘learning induced neuronal identity switch (LINIS)’. To explore the potential mechanisms underlying LINIS, we found that systemic and local (i.e. in S1) administration of the nicotinic receptor antagonist during TEC training blocked the LINIS, and concomitantly disrupted the behavior learning. Additionally, we monitored responses of two types of cortical interneurons (INs) and observed that the responses of the somatostatin-expressing (SST), but not parvalbumin-expressing (PV) INs are negatively correlated with the learning performance, suggesting that SST-INs contribute to the LINIS. Thus, we conclude that L2/3 PNs in S1 encode perceptual learning by LINIS like mechanisms, and cholinergic pathways and cortical SST interneurons are involved in the formation of LINIS.

## Introduction

Information integration in the neocortex occurs at early stages of sensory processing. During the initial phase of learning, both sensory and non-sensory information converge into the primary cortex accompanied by changes of behavioral and brain states^1–11^. Decades of work have emphasized the role of the sensory neocortex is to extract physical stimulus features whereas higher cortical areas use the sensory signals to generate decision variables, which suggest a step-by-step hierarchical processing of information in different brain regions^12–17^. Recent findings suggest that superficial layers of the sensory cortex encodes learning-related non-sensory signals, such as anticipation, attention and behavioral choice^3, 4, 11, 18–21^.

A feature of the task-related non-sensory signals’ representation is that initially quiet neurons gain the responsiveness to the sensory signals, while the neurons carried sensory signals irrespective of the behavioral outcome mostly remain unchanged^3, 4,11^. However, almost all of these findings are reward-based perceptual learning, both the reward-related and non-reward-related stimuli should be perceptive to make a correct choice. In contrast, the strategies by which the sensory cortex employs to achieve aversive associative learning remain poorly understood. Specifically, how do the neurons represent the information during learning? Does individual neurons change their responses properties at different learning stages?

To answer these questions, we simultaneously monitored animal behavior and neural activities of primary neurons (PNs) and interneurons (INs) in layer 2/3 of primary somatosensory cortex (S1) under a two-photon system throughout the learning process in the trace eyeblink conditioning (TEC) behavior learning paradigm^22–24^. Our results reveal new principals by which sensory cortices contribute to TEC learning: superficial PNs encodes learning by remain plastic and changing their response identities

## Results

### S1 is necessary for TEC acquisition but not consolidation

To investigate if and how the PNs encode the TEC-learning, we first trained the awake head-fixed mice freely moved on a running wheel to learn TEC. In our TEC paradigms, whisker deflection was the conditioned stimulus (CS, 60 Hz, 250 ms) followed by an air puff to the cornea as the unconditioned stimulus (US, 20psi, 50ms) after an interval of 250 ms (Fig. 1a). We used three different stimulation patterns in one session to isolate learning induced changes: CS paired with US (16/20 trials, CS-US), CS alone (2/20 trials, CS-only) and US alone (2/20 trials, US-only). Meanwhile, we imaged the calcium transients (via GCamP6s) of PNs through a cranial window (diameter 3mm) under a two-photon microscope after the region of interest (ROI) was identified by intrinsic imaging (Fig. 1b and Extended Data Fig. 1a, b). During imaging, mice were awake and free to walk on a 3D-printed running wheel.

**Fig. 1:**
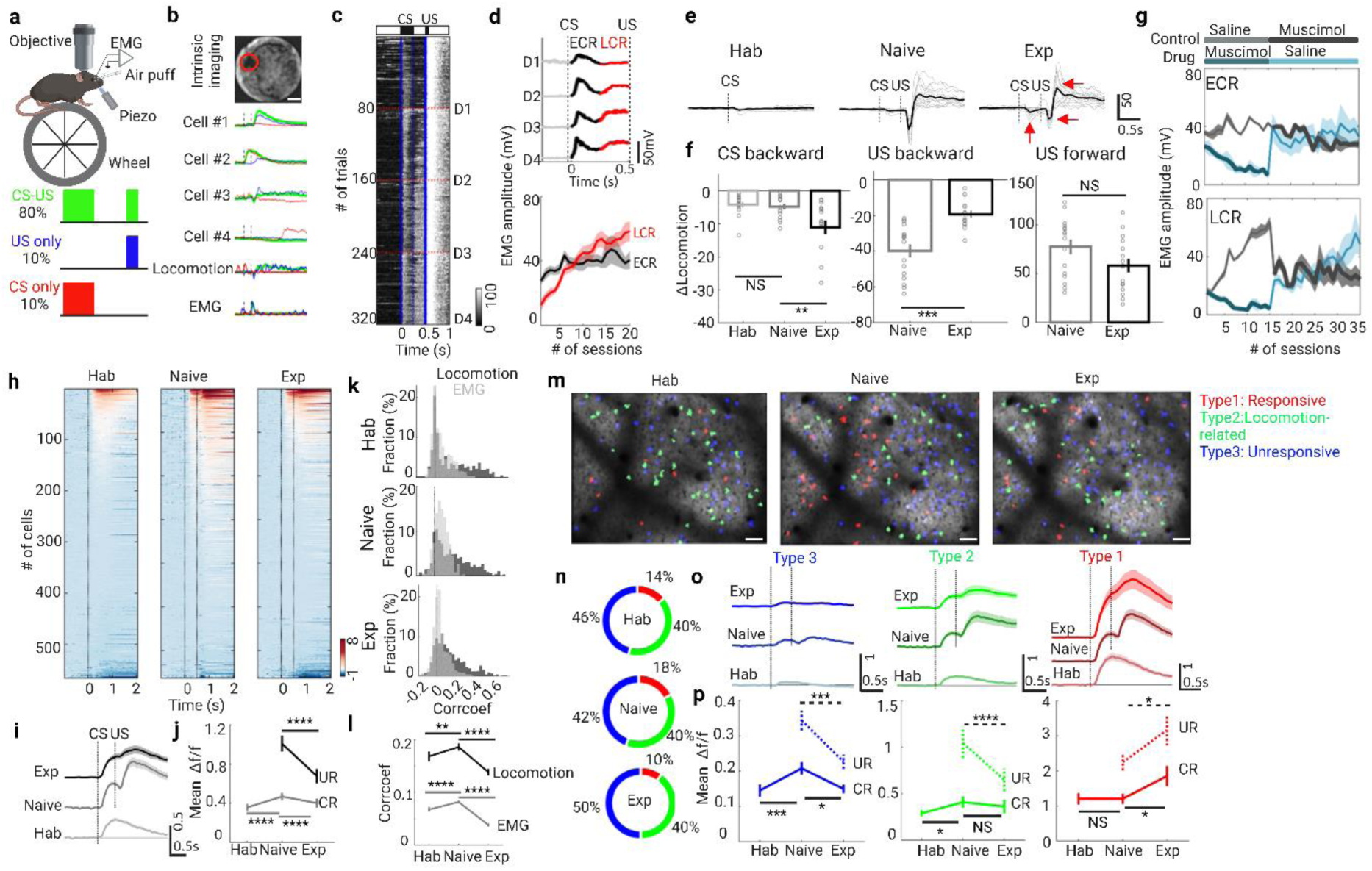
Neural population responses and behavioral patterns during TEC learning. **a**, Schematic of experimental setup (top) and trial structure (bottom). **b**, Intrinsic imaging signals are used to identify the whisker-evoked area in the barrel cortex (top, red circle, scale bar, 500mm). Average GCaMP6s signal (Δf/f) traces from four example L2/3 neurons, locomotion and EMG traces to different stimuli patterns (green, CS-US; red, CS-only; blue, US-only) in the first session of TEC training (bottom). **c**, Heat map of electromyography (EMG) responses from a representative mouse in all trials during TEC learning (16 CS-US trials in one session, five sessions in a day shown in red dashed line; blue solid lines indicate onset of CS or US, blue dashed line indicate the offset of CS). **d**, Average EMG traces of the example mouse (ECR, early conditioned response from CS onset to offset; LCR, late conditioned response from CS offset to US onset; UR not shown) in all trials during each day (top). Evolution of ECR (black) and LCR (red) during training (n = 11 mice, bottom). Data are shown as mean ± s.e.m. Shaded areas represent s.e.m. **e**, Locomotion traces change at different stages of training (Hab: habituation, only CS stimuli before TEC training; Naïve: the first session of TEC training; Exp.: expert, last session of TEC training, red arrows indicated the three patterns of locomotion: CS backward, US backward and US forward). Gray lines represent the traces from a session, black lines represent their average. The vertical dashed line indicates the stimuli onset (first one, whisker stimulation, CS; second, the air puff, US). **f**, Three parts of locomotion change differently at different stages of training. CS backward, the negative response between CS and US; US backward, the negative response after US; US forward, the maximum positive response after US (n = 15 mice; NS P=0.66, **P=0.0088, ***P=1.41 × 10^−4^, NS P=0.073; Wilcoxon test). **g**, Evolution of EMG amplitude (ECR, top; LCR, bottom) across 35 sessions of training in the control group (saline was delivered during the first 3 days and then muscimol was delivered) and drug group (muscimol was delivered during the first 3 days and then saline was delivered) (n=4 in each group). **h**, Heat maps of GCaMP6s signals (Δf/f) for all pyramidal neurons (PNs) at different stages of training (Hab, n=566 neurons from 9 mice; Naive, n=663 neurons from 9 mice; Exp, n=665 neurons from 9 mice). Signals are aligned to the CS onset. **i**, Mean signal traces of PNs at each learning stage. Data are shown as mean ± s.e.m. Shaded areas represent s.e.m. **j**, CR (gray line) and UR (black line) activity of PNs at each learning stage (CR, ****P=6.16 × 10^-^, ****P=1.43 × 10^−6^; UR, ****P=3.70 × 10^−14^; Wilcoxon test). **k**, Distribution of correlation coefficients between calcium signal and locomotion (black) / EMG (gray) at the three training stages. **l**, Correlation coefficients change at the three training stages (locomotion vs. signal, **P=0.0020, ****P=2.15 × 10^−6^; EMG vs. signal, ****P=3.96 × 10^−8^, ****P=4.16 × 10^−32^; Wilcoxon test). **m**, Two-photon fluorescence image (scale bar, 50 um) merged with different types of PNs in a representative mouse (red, responsive neurons; green, locomotion-related neurons; blue, unresponsive neurons). **n**, The fraction of 3 subtypes PNs at different training stages. **o**, Mean signal traces of 3 subtypes PNs at each learning stage. Data are shown as mean ± s.e.m. Shaded areas represent s.e.m. Data are shown as mean ± s.e.m. Shaded areas represent s.e.m. **p**, CR (solid line) and UR (dashed line) activity of 3 subtypes PNs at each learning stage (CR of others, ***P=2.33 × 10^−4^, *P=0.036; UR of others, ***P=6.73 × 10^-4^; CR of locomotion-related neurons, *P=0.015, NS P=0.061; UR of locomotion-related neurons, ****P=3.25 × 10^−5^; CR of responsive neurons, NS P=0.51, *P=0.032; UR of responsive neurons, *P=0.013; Wilcoxon test). Data are shown as mean ± s.e.m.

After repeated pairing of the CS-US over multiple sessions (6.45 ± 0.65), mice showed a gradual improvement in behavior performance, indicated by the increased orbicularis oculi muscle contractions [measured by their electromyography (EMG) amplitude (Fig. 1c)] during the time window from CS onset to US onset (CR-EMG), especially the late part of CR-EMG (LCR-EMG, Fig. 1d) but not the early part of CR-EMG (ECR-EMG) or unconditional response (UR-EMG, Extended Data Fig. 1c-h). In addition, the probability of eyeblink conditioned response (CR %) progressively increased from 11% ± 5% to 86% ± 4% (Extended Data Fig. 1c,d), which is similar to previous reports^22–24^.

Locomotion modulates the sensory processing^5, 7–10, 25^, we next investigated the characteristics of locomotion at different stages of TEC training: habituation stage, where the animals were exposed only with unreinforced CS-only before the TEC training; naive stage, which was the first reinforced session; and expert stage, which was the last session. Interestingly, we found that mice runs defensively backward both after CS and US, especially at the expert stage (Fig. 1e). We then separated the locomotion pattern into three groups: CS backward, US backward and US forward (red arrows indicated in Fig. 1e). As the learning progressed, CS backward increased while the US backward decreased (Fig. 1f). Thus, these three locomotion patterns are behavior signatures of different stages of the TEC learning.

To directly test the role of S1 in TEC learning, we inhibited neuronal activity during TEC by local infusion of muscimol (GABA_A_-receptor agonist) into S1 (Fig. 1g and Extended Data Fig. 1j). For the treatment group, muscimol was first administered then replaced by saline. For the control group the muscimol and saline were administered in the opposite order. We found that muscimol led to a reversible reduction in ECR-EMG in the treatment group, but not in the control group.

Interestingly, LCR-EMG amplitude decreased dramatically when muscimol was delivered in each group; however these differences were diminished between control and treatment group after the animals already acquired the TEC learnings, suggesting that the activities in the barrel cortex is required for TEC acquisition but not for consolidation.

### Response heterogeneity of superficial PNs across learning stages

We next examined PN activities in layer 2/3 (100 to 250 μm below the pial surface) of the S1 at different training stages. The average population activity of many PNs was modulated by the stimuli of CS or US during TEC training (Fig. 1h – j).

Surprisingly, the mean activity of the CS-triggered calcium transients (CR-signal) increased from habituation stage to naive stage (0.36 ± 0.03 vs. 0.47 ± 0.04, P=6.16 × 10^−9^) but decreased from naive stage to expert stage (0.47 ± 0.04 vs. 0.40 ± 0.04, P=1.43 × 10^−6^), which was contrary to previous findings in a different behavioral paradigms^11^. Meanwhile, the average amplitude of US-triggered calcium activity (UR-signal) decreased from naive to expert stage (1.0 ± 0.08 vs. 0.68 ± 0.07, P=3.70 × 10^−14^). Similarly, the correlation between calcium signal and locomotion or EMG displayed the same change profiles: increased from habituation stage to naive stage (locomotion: 0.17 ± 0.009 vs. 0.19 ± 0.007, P=0.0020; EMG: 0.068 ± 0.004 vs. 0.082 ± 0.003, P=3.96 × 10^−8^), but decreased from naive stage to expert stage (locomotion: 0.19 ± 0.007 vs. 0.14 ± 0.006, P=2.15 × 10^−6^; EMG: 0.082 ± 0.003 vs. 0.038 ± 0.003, P=4.16 × 10^−32^; Fig. 1k, i).

In addition to analyzing the population activity of PNs, we classified the cells into three subtypes according to their responsiveness to CS or US stimulation and locomotion (see Methods for details, Fig. 1m): responsive neurons (type 1), locomotion-related neurons (type 2) and unresponsive neurons (type 3). This classification revealed that the cellular composition changed separately across the learning stage: the fraction of responsive cells increased first from habituation to naive then decreased from naive to expert, while the fraction of type 3 displayed opposite changes and no change in locomotion-related cell populations (Fig. 1n). Additionally, the fraction of type 1 neurons was lower than 20%, while the fraction of type 3 was nearly 50%. Furthermore, the mean activity of type 1 neurons in both CS and US window increased gradually across the learning stages, while the mean activity of type 3 neurons increased first from habituation to naive and then decreased dramatically from naive to expert in both windows. For type 2 cells, the mean activity during CR and UR changed differently: it decreased during UR period dramatically from naive to expert, and it increased during CR period from habituation to naive stage only (Fig. 1o, p). Meanwhile, the mean activity of type 1 cells was far larger than that of other two subtypes, especially type 3. Based on these results, we speculate that learning induced changes of the average population activities were mainly contributed by the type 2 and type 3 cells. These results demonstrate that the response profiles of PNs are broadly different between three subsets, but also heterogeneous across learning stages.

### Learning-induced neuronal identity switch (LINIS)

Among type 1 group, responses of individual neurons were highly heterogeneous: these neurons were responsive to either CS or US (for example: cell 1 and 2 in Fig. 1b and Extended Data Fig. 1a), respectively. We next determined specific response properties of type 1 neuron by longitudinally tracking individual neurons across training sessions. To identify the same cells across sessions, we calculated the spatial coordination of each cell located in the merged vascular maps and averaged fluorescence images across the sessions (Fig. 2a; see Methods for details). The responsive cells were first detected with positive CS-evoked response (i.e. CR-neurons) at habituation stage. Then, we followed the same CR-neurons at naive and expert stages, the remainder of responsive cells were mainly responded to the US and thus they were defined as UR-neurons.

**Fig. 2:**
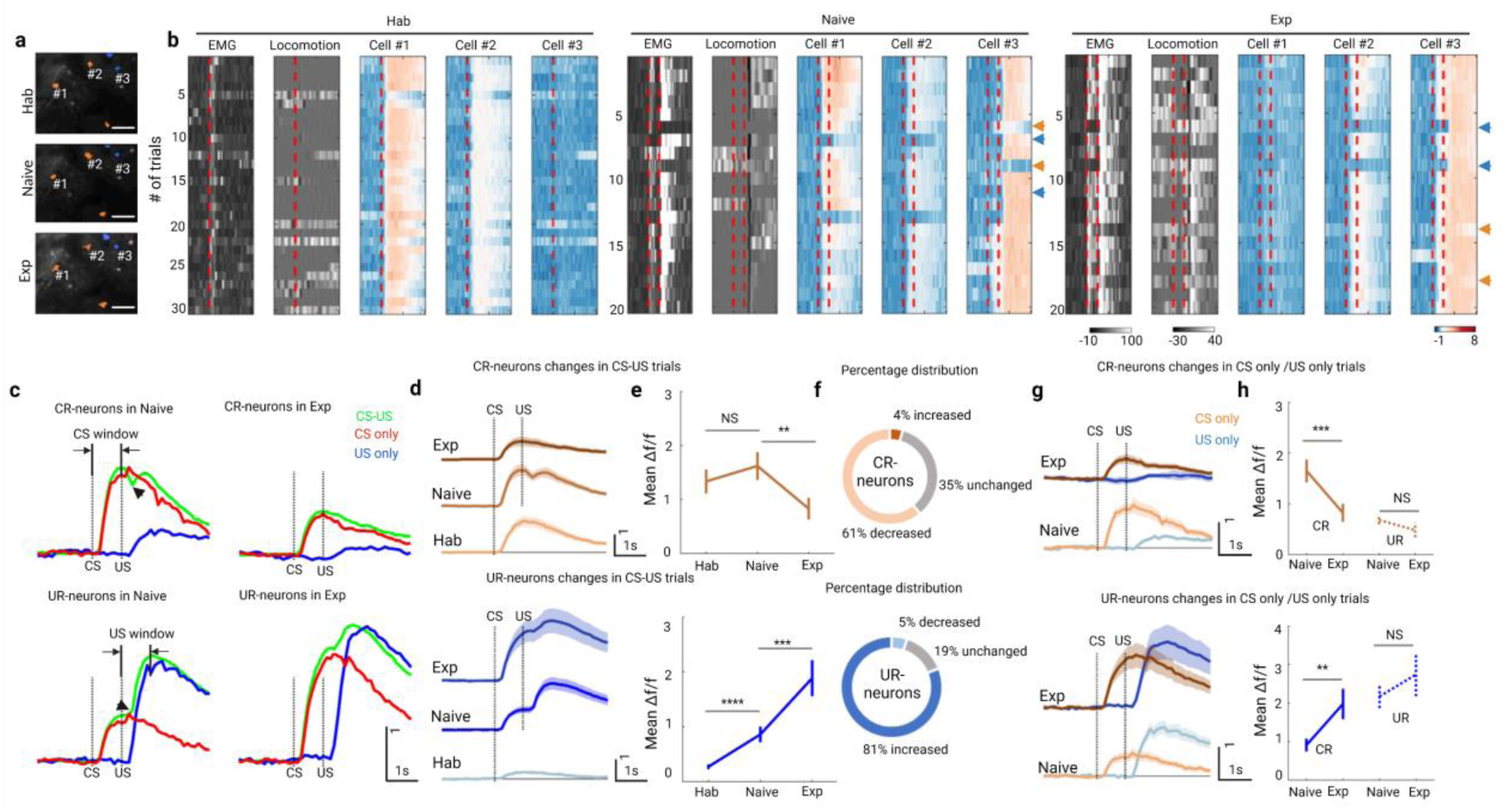
Learning induced transformation of CR– and UR– neurons. **a**, Partial field of view in layer 2/3 imaged longitudinally at each training stage (scale bar, 50 um). Longitudinally tracked neurons are colored with orange (CR-neurons) and blue (UR-neurons). **b**, Activity of the EMG, locomotion and calcium transients in each trials of the three represented neurons in A at each training stage (vertical dashed lines indicates the onset of CS or US, orange arrows indicate the CS-only trials, while blue arrows indicate the US-only trials, others are CS-US trials). **c**, Mean signal traces of CR-(top) and UR-neurons (bottom) in CS-US (green), CS-only (red) and US-only (red) trials at naïve (left) and expert (right) stage. **d**, Mean activity of CR-neurons (top) and UR-neurons (bottom) in CS-US trials at each training stage aligned to the CS onset. Data are shown as mean ± s.e.m. Shaded areas represent s.e.m. **e**, CR activity changes of CR-neurons (top) and UR-neurons (bottom) at each training stage (CR-neurons, n = 46 neurons at each stage from 7 mice; hab vs. naive, NS P=0.14; naive vs. exp, **P=0.0023; paired t-test; UR-neurons, n = 88 neurons at hab from 7 mice, n= 88 neurons at naive from 7 mice, n= 56 neurons at exp rom 7 mice; ****P=9.49 × 10^−11^, ***P=5.64 × 10^−4^; Wilcoxon test). **f**, The fraction of CR-neurons (top) and UR-neurons (bottom) at different training stages. **g**, Mean activity of CR-neurons (top) and UR-neurons (bottom) in CS-only trials (orange and brown) and US-only (blue) trials at each training stage aligned to the CS onset. Data are shown as mean ± s.e.m. Shaded areas represent s.e.m. **h**, CR activity (solid line) and UR activity (dashed line) changes of CR-neurons (top) and UR-neurons (bottom) at each training stage (CR-neurons, n = 46 neurons at each stage from 7 mice; naive vs. exp, ***P=6.97 × 10^−4^, NS P=0.20; paired t-test; UR-neurons, n= 88 neurons at naive from 7 mice, n= 56 neurons at exp rom 7 mice; **P=0.0031, NS P=0.57; Wilcoxon test).

The trial-by-trial responses of the representative CR-neurons and UR-neurons across training stages were shown in Fig. 3b. We further characterized the learning-induced dynamics of individual CR– and UR– neurons. First, decreased activities in CR-neuron and increased activities in UR-neurons occurred much earlier (from trial 14th at naive stage) than the behavioral changes (EMG and locomotion), supporting the notion that the activities of these neurons are likely to prelude behavior learning^26^.

**Fig. 3:**
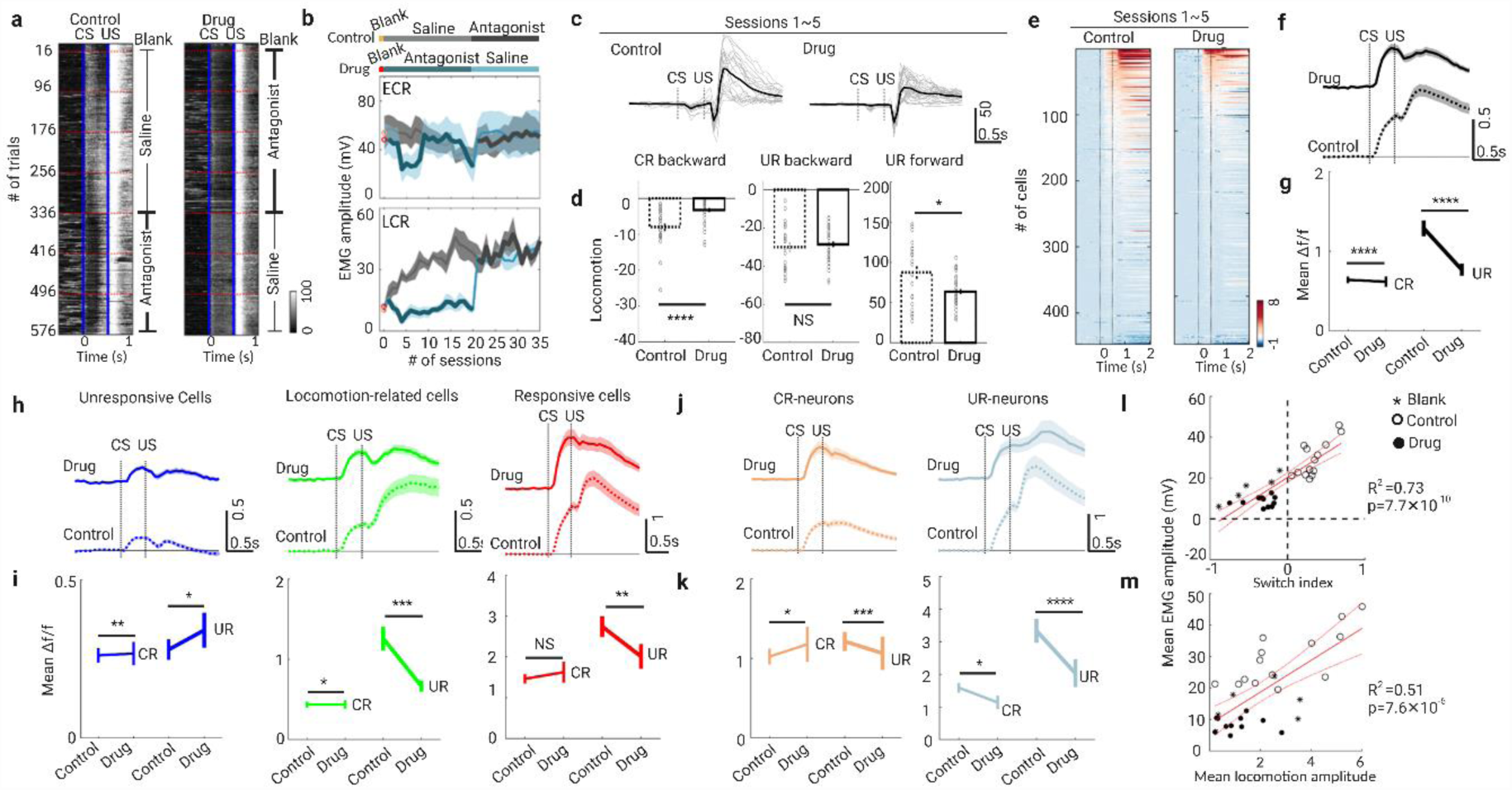
Inactivation of cholinergic signaling blocks the neuronal identity switch and impairs behavioral patterns. **a**, Heat map of EMG responses from control (left) and drug (right) mice during TEC learning (16 CS-US trials in one session, five sessions in a day shown in red dashed line; blue solid lines indicate onset of CS or US). The right side of each heat map indicates the treatment of blank, nicotinic receptor (nAChR) antagonist or saline. **b**, Evolution of EMG amplitude (ECR, top; LCR, bottom) across 35 sessions of training in the control group (One blank session in day 0, saline was delivered during the first 3 days and then nAChR antagonist was delivered) and drug group (One blank session in day 0, nAChR antagonist was delivered during the first 3 days and then saline was delivered) (n=6 in each group). Data are shown as mean ± s.e.m. Shaded areas represent s.e.m. **c**, Locomotion traces of control and drug groups during session 1∼5 of training. Gray lines represent the traces from a session, black lines represent their average. The vertical dashed line indicates the stimuli onset of CS or US. Data are shown as mean ± s.e.m. Shaded areas represent s.e.m. **d**, Three parts of locomotion change in the control and drug groups (control group, n = 23 sessions from 5 mice; drug group, n=35 sessions from 7 mice; ****P=6.6 × 10^−5^, NS P=0.52, *P=0.019, Wilcoxon test). **e**, Heat maps of GCaMP6s signals (Δf/f) for PNs in drug and control sessions 1∼5 during training (control, n=447 neurons from 3 mice; drug, n=358 neurons from 3 mice). Signals are aligned to the CS onset. **f**, Mean signal traces of PNs in drug and control sessions. Data are shown as mean ± s.e.m. Shaded areas represent s.e.m. **g**, CR (fine line) and UR (bold line) activity of PNs in drug and control sessions (CR, ****P=7.36 × 10^−5^; UR, ****P=5.56 × 10^−7^; Wilcoxon test). **h**, Mean signal traces of 3 subtypes PNs in drug and control sessions. Data are shown as mean ± s.e.m. Shaded areas represent s.e.m. **i**, CR (fine line) and UR (bold line) activity of 3 subtypes PNs in drug and control sessions (CR of unresponsive cells, **P=0.0099; UR of unresponsive cells, *P=0.013; CR of locomotion-related neurons, *P=0.012; UR of locomotion-related neurons, ***P=1.91 × 10^−4^; CR of responsive neurons, NS P=0.27; UR of responsive neurons, **P=0.019; Wilcoxon test). **j**, Mean signal traces of CR– and UR-neurons in drug and control sessions. Data are shown as mean ± s.e.m. Shaded areas represent s.e.m. **k**, CR (fine line) and UR (bold line) activity of CR– and UR-neurons in drug and control sessions (CR of CR-neurons, *P=0.021; UR of CR-neurons, ***P=5.40 × 10^−4^; CR of UR-neurons, *P=0.042; UR of UR-neuron, ****P=6.37 × 10^−12^; Wilcoxon test). **l**, Switch index versus mean EMG amplitude from CS onset to US onset in blank (asterisk), drug (solid circle) and control (hollow circle) sessions (Pearson correlation fit and 95% confidence bands, n= 31 session from 3 mice). **m**, Mean locomotion amplitude versus mean EMG amplitude from CS onset to US onset in blank (asterisk), drug (solid circle) and control (hollow circle) sessions (Pearson correlation fit and 95% confidence bands, n= 31 session from 3 mice).

Second, UR-neurons were unresponsive to CS in the habituation sessions and early trials at naive stage, but responded to CS in the later naive trials, especially during the expert sessions. Whereas CR-neurons had the opposite response profiles: responded to CS in habituation session and early trials of naive session, but decreased their responses in later trials of naive stage. These findings suggest that TEC learning mainly modulated CS-evoked responses of CR– and UR-neurons.

Third, the activity of CR-neurons within the CS window (from CS onset to US onset) in CS-US trials was almost the same as that in CS-only trials at both naive and expert stages (Fig. 2c and Extended Fig. 3a-c). Additionally, the activity of UR-neurons in the CS-US trials was the summation of that in the CS-only and US-only trials at both naive and expert stages. These results suggest that CR– and UR-neurons integrate the stimuli differently.

Additionally, as the animal gained behavioral learning, which was indicated by an increased EMG activity in CS window, CR-neurons’ activity within CS window dramatically and progressively decreased at expert stage while the UR-neurons’ activity significantly increased at both naive and expert stage (Fig. 3d, e). Meanwhile, the fraction of CR-neurons that significantly decreased their activity from naive to expert was larger than the fraction that increased (Fig. 3f; P=7.36×10^−9^, Chi-square test). In contrast, the percentage of UR-neurons whose activity increased were larger than that of UR-neurons whose activities were decreased (Fig. 3f; P=7.67×10^−16^, Chi-square test).

Lastly, CR-neurons’ activity in CS-only trials was larger than that in US-only trials, and decreased dramatically from naive to expert stage (Fig. 3g,h). In contrast, UR-neurons’ activity in CS-only trials was lower than that in US-only trials, and increased significantly from naive to expert (Fig. 3g,h). Meanwhile, activity of both CR– and UR-neurons in US-only trials has no significant changes from naive to expert. In summary, learning induced CR-neurons responding to the whisker stimuli lose their ability of sensory perception, however, the previously unresponsive UR-neurons gain responsiveness to whisker stimuli, a novel cellular phenotype we described here as ‘learning induced identity switch (LINIS)’.

### Cholinergic signaling mediates LINIS and behavioral learning

What is the mechanism underlying LINIS? Previous works have supported a role of acetylcholine (ACh) in mediating learning-related modulations of neural activity^27–29^. To test whether the cholinergic system modulates the PNs’ activity during TEC learning, we first pharmacologically manipulated cholinergic signaling systematically while imaging the calcium transients of PNs across learning stages. The strategy of drug delivery was as following (Fig. 3a, b and Extend Fig 4.a): after habituation and the first session of training on day 1 (session 0), nicotinic receptor antagonist (mecamylamine, 5 mg kg^−1^) was delivered systemically once a day 10 minutes before training (session 1 to session 20) from day 2 to day 5, the antagonist was replaced with saline from day 6 to day 8. In the control group, the saline was first injected from day 2 to day 5, then it was replaced with the antagonist from day 6 to day 8.

**Fig. 4:**
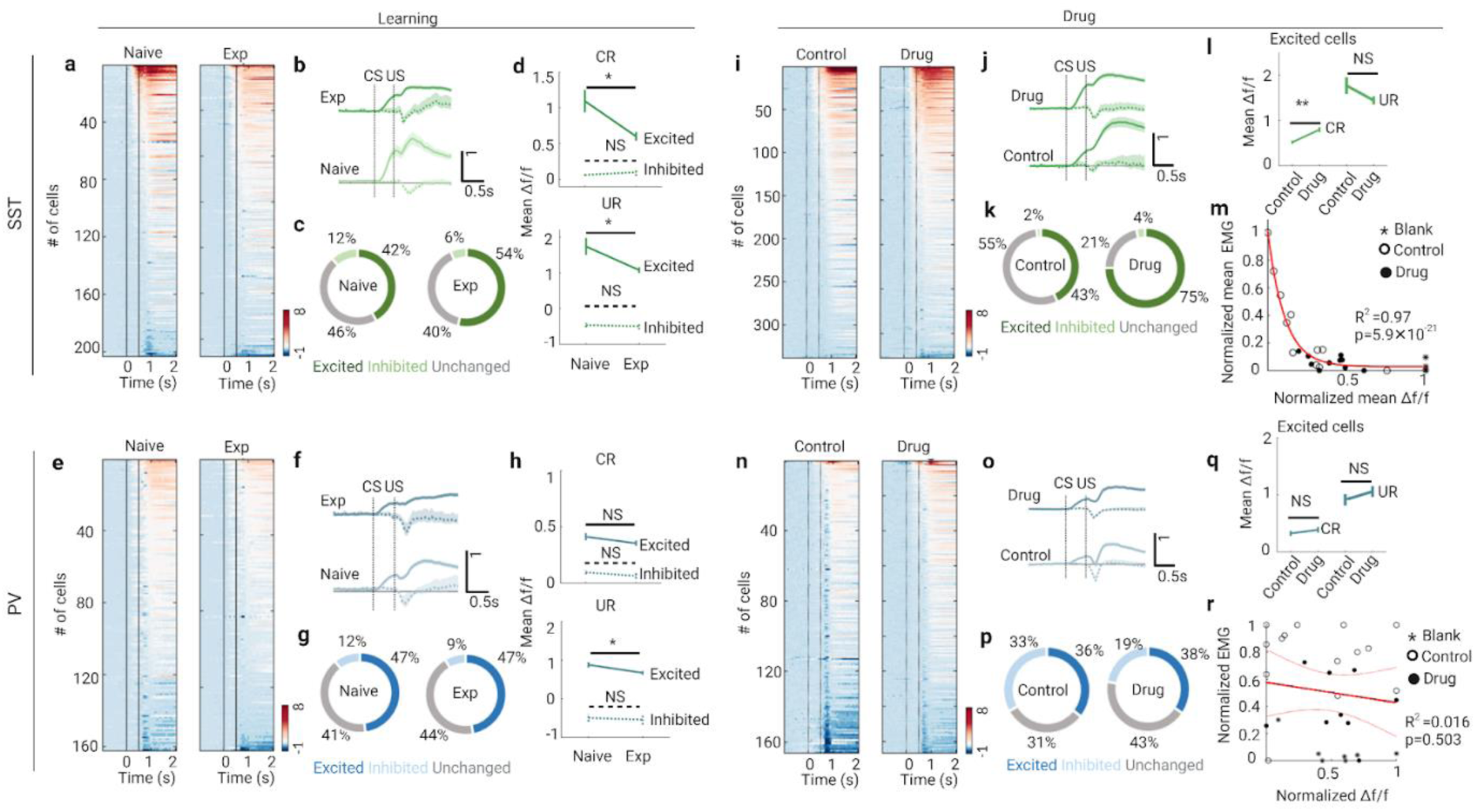
Activities of SST-INs, not PV-INs are negatively correlated with the learning performance. **a**, Heat maps of GCaMP6s signals (Δf/f) for SST-INs in CS-US trials at different stages of training (Naive, n=203 neurons from 7 mice; Exp, n=255 neurons from 7 mice). Signals are aligned to the CS onset. **b**, Mean signal traces of SST-INs in CS-US trials at each learning stage (solid line indicates the excited cells, dashed line indicates the inhibited cells). Data are shown as mean ± s.e.m. Shaded areas represent s.e.m. **c**, The fraction of excited and inhibited SST-INs at different training stages. **d**, CR (top) and UR (bottom) activity of excited (solid line) and inhibited (dashed line) SST-INs at different training stages (CR of excited cells, *P=0.016; CR of inhibited cells, NS P=0.53; UR of excited cells, *P=0.033; UR of inhibited cells, NS P=0.92; Wilcoxon test). **e**, Heat maps of GCaMP6s signals (Δf/f) for PV-INs in CS-US trials at different stages of training (Naive, n=162 neurons from 5 mice; Exp, n=153 neurons from 5 mice). Signals are aligned to the CS onset. **f**, Mean signal traces of PV-INs in CS-US trials at each learning stage (solid line indicates the excited cells, dashed line indicates the inhibited cells). Data are shown as mean ± s.e.m. Shaded areas represent s.e.m. **g**, The fraction of excited and inhibited PV-INs at different training stages. **h**, CR (top) and UR (bottom) activity of excited (solid line) and inhibited (dashed line) PV-INs at different training stages (CR of excited cells, NS P=0.43; CR of inhibited cells, NS P=0.28; UR of excited cells, *P=0.010; UR of inhibited cells, NS P=0.75; Wilcoxon test). **i**, Heat maps of GCaMP6s signals (Δf/f) for SST-INs in drug and control sessions 1∼5 during training (control, n=338 neurons from 3 mice; drug, n=327 neurons from 3 mice). Signals are aligned to the CS onset. **j**, Mean signal traces of excited (solid line) and inhibited (dashed line) SST-INs in drug and control sessions. Data are shown as mean ± s.e.m. Shaded areas represent s.e.m. **k**, The fraction of excited and inhibited SST-INs in drug and control sessions. **l**, CR (fine line) and UR (bold line) activity of excited SST-INs in drug and control sessions (CR, **P=0.0013; UR, NS P=0.13; Wilcoxon test). **m**, Normalized mean SST-INs’ signal versus normalized mean EMG amplitude from CS onset to US onset in blank (asterisk), drug (solid circle) and control (hollow circle) sessions (decay function fitting, n= 30 session from 3 mice). **n**, Heat maps of GCaMP6s signals (Δf/f) for PV-INs in drug and control sessions 1∼5 during training (control, n=166 neurons from 2 mice; drug, n=333 neurons from 3 mice). Signals are aligned to the CS onset. **o**, Mean signal traces of excited (solid line) and inhibited (dashed line) PV-INs in drug and control sessions. Data are shown as mean ± s.e.m. Shaded areas represent s.e.m. **p**, The fraction of excited and inhibited PV-INs in drug and control sessions. **q**, CR (fine line) and UR (bold line) activity of excited PV-INs in drug and control sessions (CR, NS P=0.57; UR, NS P=0.32; Wilcoxon test). **r**, Normalized mean PV-INs’ signal versus normalized mean EMG amplitude from CS onset to US onset in blank (asterisk), drug (solid circle) and control (hollow circle) sessions (Pearson correlation fit and 95% confidence bands, n= 31 session from 3 mice).

In the mecamylamine-treated group (drug group), LCR-EMG remained at a low amplitude until the mecamylamine was replaced with saline. In the control group, LCR-EMG increased gradually with progression of training, and no significant changes was found when the saline was changed to mecamylamine. Meanwhile, no significant difference was observed in ECR-EMG across or within the two groups. To confirm the effect of nAChRs on the LCR-EMG, we next locally delivered the mecamylamine to the S1 region with an implanted cannula (Extend Fig 4.b). Similarly to the effects of systemic administration of mecamylamine, only LCR decreased significantly in the sessions treated with mecamylamine. These findings demonstrate that only LCR which correlated with the learning performance was modulated by the nicotinic receptors, acting locally at S1.

We next examined the roles of muscarinic AChRs in learning by applying muscarinic antagonist (atropine, 5 mg kg^−1^) in separate groups of animals (Extend Fig 4.c). In contrast to the effects of nicotinic receptor antagonist, atropine did not robustly impaired the learning performance. Additionally, optogenetic activation of the cholinergic fibers (via ChAT-ChR2 mice) in S1 increased the LCR during TEC training (see Methods for details, Extend Fig 4.d, e). These manipulations provide direct evidence that cholinergic activity in S1 is crucial for the learning performance. Concurrent with the alteration of LCR, cholinergic signaling antagonism also impaired the stereotypical locomotion patterns, resulting in a reduced amplitude of backward locomotion in the CS window and reduced amplitude of the forward locomotion in the US window (Fig 3.c, d). However, no significant changes were found in the amplitude of backward locomotion after US onset.

Lastly, inactivating cholinergic signaling with nAChR antagonist (mecamylamine) dramatically decreased the average activity of population PNs in both CS-induced responses (CR) and US-induced responses (UR), especially the signal amplitude of UR (Fig. 3e-g). We further analyzed the activities of the three subsets of PNs (Fig. 3h,i): responsive cells (type 1), locomotion-related cells (type 2) and unresponsive cells (type 3). We found that CR in type 2 and type 3 increased but UR in these two subtypes decreased dramatically in drug sessions compared with saline-treated sessions. For the activity of responsive neurons (type 1), only decreased UR in drug sessions were observed. For type 2 cells (i.e. responsive cells), we furtherly measured the activity of CR-neurons and UR-neurons separately (Fig. 3j, k). We found the increased CS-induced responses in CR-neurons and decreased CS-induced responses in UR-neurons after treated with mecamylamine, which may result in unchanged CS-induced responses of all responsive cells. Both CR-neurons and UR-neurons in drug sessions showed opposite dynamics when compared with that from naive to expert sessions. Therefore, we speculated that the activity difference in CR-neurons and UR-neurons may be the key for TEC learning performance. The switch index (SI, see Methods for details) measuring the absolute difference in calcium transients within the CS window of CR-neurons and UR-neurons was calculated. Notably, the SI in each session correlated positively with the mean LCR-EMG amplitude in each session (Fig. 3l). Also, the mean amplitude of forward locomotion (absolute value) showed a high positive linear relationship with the mean amplitude of LCR-EMG in each session (Fig. 3m). These findings suggest that the LINIS encodes the CS and US efficiently during TEC learning, which is modulated by the cholinergic pathways.

### SST-INs, but not PV-INs contribute to the LINIS

Accumulating evidence suggested that different cortical interneurons (INs) play distinct roles in learning^6, 27, 30–32^. To investigate whether (and which subtype of) INs contributed to the TEC learning, we monitored longitudinally the calcium transients of somatostatin-expressing (SST) and parvalbumin-expressing (PV) INs during training.

According to the response profile to the CS and US, we categorized the INS into three subtypes (see Methods for details): excited, inhibited and unchanged cells. Interestingly, the inhibited INs of both SST and PV responded only to the US but not CS, whereas the excited cells responded to both CS and US (Fig. 4a, b, e, f). We found that the fraction of excited SST-INs or PV-INs was significantly larger than that of inhibited INs (Fig. 4c; SST-INs at naive stage P=9.76×10^−12^, SST-INs at expert stage P=3.26×10^−16^; Fig. 4g; PV-INs at naive stage P=1.44×10^−5^, PV-INs at expert stage P=2.22×10^−16^; chi-square test). Across different learning stages, activities of excited SST-INs decreased dramatically in both ‘CS-window’ (CR) and ‘US-window’ (UR), while that of the inhibited SST-INs were not modulated by learning (Fig. 4d).

For PV-INs, only the activity of inhibited subtype in the ‘US-window’ decreased at expert stage (Fig. 4h). These findings indicate that only population activities of the excited SST-INs code the identities of CS and US stimuli. To investigate whether activity of INs was also modulated by cholinergic signaling during TEC learning, we performed pharmacological blockade of cholinergic signaling systematically while imaging the calcium transients of INs across the learning stages. The strategy of drug delivery in Ai96/SST or Ai96/PV mice was the same as that in Ai96/CamKII mice.

Inactivating cholinergic signaling with nAChR antagonist (mecamylamine, 5 mg kg^−1^) dramatically increased the average activity of excited SST-INs but not inhibited SST-INs in the ‘CS-window’ but not in the ‘US-window’ (Fig. 4i-l and Extended Fig. 5a-h). Notably, the mean excited SST-INs’ activity in each session correlated negatively with the mean LCR-EMG amplitude in each session (Fig. 4m). Repeating the same analyses on the PV-INs, inhibited PV-INs but no excited PV-INs decreased their CR activities but increased the UR activity in the mecamylamine-treated sessions (Fig. 4e-r and Extended Fig. 5i-p). These results confirm a vital role of excited SST-INs during TEC learning. Together, these findings suggest that excited SST-INs are negatively correlated with the learning performance, which support the role of SST-INs in the formation of LINIS in PNs.

## Discussion

We trained mice to perform a TEC task in which the mice learned to close the eyelid before the delivery of air puff. We quantified the performance of the task by orbicularis oculi muscle EMG and arousal state by locomotion, as well as the activities of single excitatory and inhibitory neurons in the somatosensory cortex. In addition, we evaluated the relationship between neuronal responses and behavior by pharmacological manipulating the nicotinic receptors. We found that: (i) the heterogeneous neuronal representations of both the modality-specific sensory stimuli and non-specific stimuli in barrel cortex exhibit distinct properties at different learning stage; (ii) both the direction and amplitude of locomotion modulated by learning performance and occurred prior to the appearance of task-relevant stimuli; (iii) a small populations (∼20%) of PNs but with distinct response properties, i.e., the sensory-specific neurons (CR-neurons) and behavioral-sensitive neurons (UR-neurons), were transformed oppositely with learning (here we coined ‘LINIS’); (iv) behavioral performance was positively correlated with LINIS and negatively correlated with SST-INs activity, which were modulated by cholinergic signaling. Our results pinpoint a crucial role of the primary sensory cortex in Pavlovian learning and provide evidence for a new strategy of sensory cortex for sensorimotor learning via LINIS like mechanisms.

### LINIS is an efficient strategy for TEC learning

Sensory cortex receives inputs from multiple sources carrying a variety of information and needs to establish effective representations of behaviorally important stimuli^33^. We observed that the locomotion-related neurons were separated from the responsive neurons. Meanwhile, as the training progress, a division of labor takes place within overall population of PNs such that they showed distinct characteristics at different learning stages. Additionally, INs of L2/3 in S1 also showed division of labor as well. Majority of SST-INs and PV-INs were excited by CS or US, while a small proportion were inhibited by US and did not respond to the CS. However, SST-INs but not PV-INs were involved in learning, which is consistent with the previous findings^2, 27, 32^.

Importantly, we found a small populations of PNs (∼20%) with distinct properties: CR-neurons (mainly responsive to CS) and UR-neurons (mainly responsive to US), were transformed oppositely by the TEC learning. CR-neurons decreased their responsiveness to whisker stimulation (i.e. CS), while UR-neurons increased their responses to whisker stimulation (here we called ‘learning-induced identity switch’, LINIS). However, earlier studies observed that the responsiveness to the sensory stimulus mostly remained unchanged during learning, which is different from the CR-neurons’ response pattern here^4, 11^. We speculate the reason for this difference is mainly from the different behavioral paradigms, especially the reward versus punishment and/or freezing versus running^34–36^. In some perceptual learning paradigms^3, 11^, the mouse received two different stimuli, for example, vertical and horizontal visual stimulation; but in our TEC paradigm, a single CS (i.e. 60 Hz whisker stimuli) was used.

UR-neurons’ response pattern during learning acquisition were similar to the studies reported earlier^3, 4, 11^, where learning recruits previously unresponsive neurons to enhance their responses was consistent with. The response profiles of UR-neurons in our study are almost the same as the task-relevant neurons in the perceptual learning, which showed enhanced and preceded activity during learning.

Additionally, the CS-induced response in UR-neurons would disappear when there are no whisker stimuli, and the activity in CS-US trials of UR-neurons was the summation of that in CS-only and US-only trials. These results suggest that UR-neurons have the ability to sum the sensory-specific and non-specific signals, which provide another example about the changes of value-sensitive neurons in sensory cortex resulting from the learning-induced new stimuli-like signal.

### Cholinergic signaling modulates LINIS via SST-INs

The running speed of the animal is often used as measures of the arousal state^7, 25, 37, 38^. The increased locomotion after delivery of air puff suggested an increase in the arousal state. Importantly, we found both the speed and direction of locomotion varied differently in different time windows according to the temporal characteristic of stimuli delivered: CS window, early and late US window. Additionally, they were modulated separately by learning, which indicates that three patterns of locomotion may originated differentially. ‘US-backward’ is likely to be the startle response: it was a defensive running away behavior. It was reduced at the expert stage, presumably due to animals are familiar with the US and thus reduced fear. The ‘CS-backward’ may reflect the defensive motor planning to avoid the incoming harmful US, based on the premotor theories^39, 40^.

Higher arousal is associated with spatiotemporally dynamic patterns of cholinergic modulation in sensory cortex^41^. Cholinergic signaling from the basal forebrain (BF) could rapidly convey time-locked information to the sensory cortex about the behavioral state of the animal and the occurrence of salient sensory stimuli^29, 42^. We demonstrated that inactivation of nAChRs decreased the ‘CS-backward’ and ‘US-forward’ locomotion, but had no effects on the ‘US-backward’, which further confirmed the difference between the ‘CS-backward’ and ‘US-backward’ locomotion. Also, inactivation of nAChRs led to the loss of neuronal activity after air puff, which suggests that both locomotion and neuronal responses were modulated by the cholinergic signaling. In particular, we found that the CR activity of CR-neurons increased, whereas CR activity of UR-neurons decreased after treated with nAChRs antagonist, which suggests that the salience of information relayed to downstream areas was disrupted by the antagonist. The high linear relationship between the switch index (which is the measurement of LINIS) and learning performance suggested that the discriminability of S1 came from the comparison of responses from CR-neurons and UR-neurons.

Additionally, electrical stimulation of the BF revealed acetylcholine could rapidly modulate the neural activity in the visual cortex, importantly, both the incidence and the polarity of these responses depended on cell-type^42, 43^. The majority of the targeted neurons are INs but not PNs^42, 44, 45^, especially only muscarinic receptors are widely expressed on PNs^46, 47^. Our results in S1 showed that blocking the muscarinic receptors had no significant influence on the learning performance. Therefore, the PNs’ activities may be modulated indirectly by the cholinergic signaling during learning.

Increasing evidence supports that feedback from high-order brain regions act through INs during learning^2, 6, 27, 48–51^. Our results support that both SST-INs and PV-INs were regulated by nAChRs inhibition, and the subtypes of these INs were differentially modulated by nAChRs. Furthermore, we confirmed the negative relationship between excited SST-INs activity and learning performance, which suggests that the increased UR-neurons’ activity may results from the decreased response of SST-INs, but not PV-INs. However, we have not found the increased responsiveness of INs subtypes during learning, which suggest the learning-induced decreases of CR-neurons were not modulated by the INs but may result from the modulation of higher order brain regions, such as retrosplenial cortex^1^ and posterior medial (POm) nucleus of the thalamus^2, 49, 52^.

In summary, we conclude that L2/3 PNs in S1 encode discriminability efficiently by LINIS like mechanisms, and cholinergic pathways and cortical interneurons are involved in the formation of LINIS.

## Methods

### Subjects Details

All procedures were approved by the Institutional Animal Care and Use Committee (IACUC) and the Biosafety Committee of the University of Wyoming^53, 54^. Immune-competent mice of both sexes (age range p60–p300) were used in the experiments described here. Food and water were available *ad libitum*. Mice were first housed in groups of 2∼5 and after surgery were housed alone in a vivarium maintained at 21-23°C on a 12 hr light/dark cycle. All imaged mice were derived from the crosses of Ai96 [B6;129S-Gt(ROSA)26Sortm96.1(CAG-GCaMP6s)Hze/J (JAX Stock No: 024106)] with CamKII-Cre (JAX Stock No: 005359), PV-Cre (JAX Stock No: 008069) or SST-cre (JAX Stock No:013044). For optogenetics, ChAT-ChR2 (JAX Stock No: 014546) mice were used. Mice expressing the transgenes (Ai96 and cre-specific) or ChAT-ChR2 were identified by PCR, outsourced to Transnetyx (transnetyx.com).

### Surgeries

Mice were anesthetized with 3% isoflurane (v/v) and maintained with 0.4 LPM oxygenated 2% isoflurane (vol/vol) throughout the surgery^53, 54^. The animals were mounted on a stereotaxic device (NARISHIGE SG-4N) and maintained at 37°C using a heat pad (K&H no.1060). 70% isopropyl alcohol and iodine were placed on the incision site. The scalp was shaved and cleaned, and the exposed skull was allowed to dry. Then the exposed skull and wound margins were covered by a thin layer of Vetbond. Once dry, a thin layer of dental acrylic was applied around the wound margins, avoiding the region of skull overlying S1 (right hemisphere, 1.6mm posterior and 3.5mm lateral of bregma). A metal head bar was affixed with dental cement caudally to the skull. After implantation, the remainder of the exposed skull were sealed with dental cement. Mice were then recovered on a heating pad. When alert, they were placed back in their home cage. Carprofen was administered daily for 3 days post-surgery. Mice were left to recover for at least 7 days prior to task training.

For recording the differential electromyography (EMG) activity of the ipsilateral orbicularis oculi (OO) muscle, two polyimide-insulated stainless steel wires (125μm, California Fine Wire) and connecting pins were implanted to the upper eyelid of the left eye^23^. The third wire was implanted to the skull near the olfactory bulb as a ground.

For two-photon imaging^55, 56^, a 3.5 mm circular piece of skull was removed above S1 (right hemisphere, 1.6 mm posterior and 3.5 mm lateral relative to bregma). Once the skull was removed, a sterile 3 mm diameter cover glass was placed directly on the exposed dura and sealed to the surrounding skull with Vetbond.

For in vivo cannula infusion^57^, a 1 mm circle craniotomy was drilled above S1 (right hemisphere, 1.6 mm posterior and 3.5 mm lateral relative to bregma).Then a guide cannula (2 mm; C315GMN/SPC; Bilaney) was implanted. The cannula was secured with Vetbond.

For optogenetics^57^, a 1 mm circle craniotomy was drilled above S1 (right hemisphere, 1.6 mm posterior and 3.5 mm lateral relative to bregma). Then an optic fiber (MFC_400/480-0.22-0.5MM_MF1.25_FLT, Doric Lenses, Quebec, Canada) was implanted and secured with Vetbond.

### Data Acquisition

For intrinsic signal imaging^4, 58^, the C2 barrel column was located using intrinsic signal imaging under anesthesia. The individual whisker (C2) was inserted into a capillary connected to a piezoelectric device at 8 Hz (1s, with a 10-s gap between pulse trains) controlled by PulsePal via Matlab. A red light source to illuminate the surface through the cranial window. Reflectance images 300 µm below the brain surface were required with ×2.7 objective. Vasculature image was then acquired by a 546-nm interference filter, and superimposed on the intrinsic signal image.

For two-photon imaging^55, 56^, two-photon calcium imaging was achieved using a resonant/galvo scanning two-photon microscope (Neurolabware, Los Angeles, CA) controlled by Scanbox software (Los Angeles, CA). GCaMP6s was excited by an Insight X3 laser (Spectra-physics, Milpitas, USA) setting to 920 nm focused through a 16×/0.8NA water-immersion objective. Images (512 x 796 pixels, 490×630 μm) were acquired with 15.5 Hz frame rate at a depth of 120 to 320 μm below the pial surface for layer 2/3 imaging. During acquisition, the mouse moved free on a 3D-printed running wheel. The locomotion was recorded by a rotary encoder with its post through the wheel axle. The rotary encoder was triggered at scanning frame rate. And the TTLs of the whisker stimulation and air puff were output via the PulsePal to the two-photon system which were used to align the image

For EMG acquisition^23, 59^, the eyelid movement signal was band-pass filtered with 50–500 Hz and amplified using a 4-channel differential amplifier (model 1700, A-M Systems) and acquired with the 16-bit data acquisition system (Digidata 1322A, Axon Instrument) through an analog input channel sampled at 1000 Hz. And the TTLs of the whisker stimulation and air puff were output via the PulsePal to the data acquisition system which were used to align the EMG signal.

To deliver the whisker stimulation, E-650 Piezo Amplifier and piezo controlled by PulsePul were used. A solenoid valve (PSV-5, Aalborg) and the driver controlled by PulsePal were used to deliver the air puff to the eye. The control of the two-photon system running and delivery of whisker stimulation, air puff and laser stimuli was controlled by custom-made software in Matlab (MathWorks).

### Behavioral Procedures

Prior to training, mice were first habituated to head restraint for 3 d by placing them on the wheel with their heads fixed for 1 h. From 4th day to 6th day, whisker stimuli (10Hz) was delivered in 4 sessions (30 trials per session) and meanwhile, two-photon imaging was conducted according to the results of intrinsic signal imaging^22,24^. On the 7th day, the whisker stimuli was changed to 60 Hz but in 1 session (30 trials), which was named as ‘Habituation stage during learning (Hab)’ in this article.

After habituation, trace eyeblink conditioning (TEC) was carried out. The conditional stimulus (CS) was whisker deflection (60 Hz, 250 ms in duration) followed by unconditioned stimulus (US) after an interval (250 ms from CS offset to US onset). The US was a periorbital airpuff (15 psi, 50 ms in duration, delivered via a plastic needle placed 5 mm from the cornea and pointed at it). The interval between trials varied from 20 s to 40 s randomly. Each mouse was trained in 5 consecutive sessions per day (20 trials per session) till the mouse learned to maintain the closure of eyelids from the CS onset to the US offset (which was reflected by the EMG processing. Details in data analysis). The first training session was named ‘Naive stage during learning (Naive)’ and the last training session was named ‘Expert stage during learning (Exp)’. The stimuli of 20 trials in one session were also randomly presented, with a proportion 80% paired of CS and US (CS-US), 10% CS alone (CS-only) and 10% US alone (US-only). If a mouse showed signs of discomfort, it was returned to the home cage for up to 24 h.

### Pharmacological and optogenetic manipulation of cholinergic inputs

All drugs or saline were delivered starting from the second session of training. The drug or saline were injected at the beginning of each day. After waiting 10 minutes, the TEC training continued (5 sessions per day). All drugs were dissolved in physiological saline (0.9% sodium chloride). For systemic inactivation of cholinergic inputs, nicotinic acetylcholine receptor antagonist mecamylamine (5 mg kg^−1^,M9020, Sigma-Aldrich) and muscarinic acetylcholine receptor antagonist atropine (5 mg kg^−1^, Y000878, Sigma-Aldrich) were injected intraperitoneally (i,p.)^27, 60^. For local inactivation, mecamylamine (0.5 ug/ul) and muscimol (0.1ug/ul, M1523, Sigma-Aldrich) were delivered through the implanted cannula. Pressure injection using a customized device driven by a single axis hydraulic manipulator (NARISHIGE mmo-220A) at a rate of 30-50 nL/min.

For optogenetic activation, the blue light (M470F3, Thorlabs) was delivered from the CS offset to the US onset through the implanted optic fiber and controlled by the LED-driver (LEDD1B, Thorlabs).

### Data processing and analysis

The pipeline for the two-photon images was processed as previously described^55, 56^. Briefly, movies from the same plane for each session were concatenated and motion corrected. After alignment, the mean image generated was used to track the same cells for longitudinal imaging. For segmentation, a MATLAB graphical user interface tool (Scanbox, Los Angeles, CA) was employed to define regions of interest (ROI) corresponding to neuronal soma. After segmentation, the calcium transient of each cell was extracted and estimated spiking was calculated via non-negative temporal deconvolution of the corrected ROI signal using Vanilla algorithm.

For calcium transients, we first calculated the baseline of the signal (∼930 frames, 60s duration before CS onset) in each cell by choosing the minimum of the mean values in the two parts (1∼450 frames and 451∼900 frames). Then the ratio of Δf/f at each time point was calculated. During each session, the signal was segmented into 30 or 20 trials (30 trials at Hab stage, 20 trials at Naive and Exp) aligned by the onset of CS (1s before CS onset and 2s after CS onset).

For cell responses classification, PNs were classified as ‘responsive’, ‘locomotion-related’ and ‘others’ cells based on the responses of the cells. The responsive neurons were defined by the trial number of estimated spikes after CS onset was larger than 10 (60% of the CS-US trials) in CS-US trials. The locomotion-related cells were determined by the correlation coefficient of the cell was larger than 8th percentile of all of cells’ correlation coefficient distribution in one session. To avoid contamination of the stimulation evoked responses, the evoked signals/locomotion (10s time window after CS onset) was removed from the session. The overlap of the responsive neurons and locomotion-related cells were defined as locomotion-related cells. The remainder of the cells are ‘unresponsive cells‘. Responsive neurons were divided into two types: CS responsive neurons (CR-neurons) and US responsive neurons (UR-neurons) based on the maximum amplitude comparison during CS window (500 ms duration from CS onset to US onset) and US window (500 ms duration after US onset).

For longitudinal tracking of the same cells, the same imaging plane was located across sessions using images acquired previously as reference by the vascular map and averaged fluorescence images. After imaging, we merged the vascular map and two-photon image in each session, then we chose one same cell in all merged images as a reference cell. By calculating the spatial locations to the reference cell, we could get the same cells across sessions.

INs (SST– or PV-INs) were classified as ‘excited’, ‘inhibited’ and ‘unchanged’ cells based on their responses to CS. The excited cells were defined when the maxima of the mean signal (absolute value, value 1) from 16 CS-US trials 0∼2s after the CS onset were above 4 × s.d. of the baseline. The inhibited cells defined when the minimum of the mean signal (absolute value, value 2) from 16 CS-US trials 0∼2s after the CS onset were above 4 × s.d. of the baseline. The overlap of the excited cells and inhibited cells were distinguished by the comparison of the value 1 and 2 (excited: value 1> value 2; inhibited value 1<= value 2). The remainder of the cells are ‘unchanged cells‘.

For locomotion analysis, we first calculated the baseline of the locomotion (∼930 data, 60s duration before CS onset) by choosing the minimum of the mean values in the two parts (1∼450 frames and 451∼900 frames). Then the baseline was subtracted. During each session, the locomotion traces were segmented into 30 or 20 trials (30 trials at Hab stage, 20 trials at Naive and Exp) aligned by the onset of CS (1s before CS onset and 2s after CS onset).

For EMG analysis^23, 59^, we first aligned the length of EMG same to locomotion or calcium transient by resampling the EMG original data according to the TTLs of CS and US. Then, we calculated the baseline of the EMG (∼930 data, 60s duration before CS onset) by choosing the minimum of the mean values in the two parts (1∼450 frames and 451∼900 frames). Then the baseline was subtracted. During each session, the EMG traces were segmented into 30 or 20 trials (30 trials at Hab stage, 20 trials at Naive and Exp.) aligned by the onset of CS (1s before CS onset and 2s after CS onset). CS response (CR) in each trial was defined as the EMG activity 4 s.d. above the baseline, present at least 200 ms before US onset. CR percentage (CR %) within a session larger than 60% was considered to reach the learning criterion. The early CS response (ECR) was the mean amplitude of EMG from CS onset to CS offset, while the late CS response (LCR) was the average amplitude of EMG from CS offset to US onset. The US response (UR) was the mean amplitude of EMG after US onset (500 ms duration after US onset).

For linear regression, the switch index was calculated by the equation:

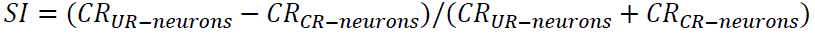

Then linear regression was fitted (Matlab ‘fitlm’ function). For decay function fitting, the equation was:

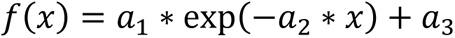

Then the Matlab function (‘fitnlm’) was used for fitting the data.

### Statistical analysis

All the statistical details are indicated where used. All of the data were expressed as the mean ± standard error of the mean (s.e.m.). Statistical analyses were conducted using Matlab. Parametric tests were used whenever possible to test differences between two or more means. Non-parametric tests were used when data distributions were non-normal. For pairwise comparisons of proportions, Chi-square test was used. Two-tailed tests were used unless noted otherwise. A value of P < 0.05 was considered to be statistically significant.

## Acknowledgements

We thank Dr. Z. Zhang for imaging and C. Zhang for animal husbandry, histology assistance and items purchasing. We thank Dr. Y. Li and Y. Wang for discussion. This work is supported by grants from National Institute of Mental Health (1R21MH131363-01) and from National Institute of General Medical Sciences (2P20GM121310).

## Author contributions

Q.S. designed the experiment and supervised the project, acquired the funding, and edited the manuscript. J.D. built the behavioral set-up, co-designed the experiments, collected and analyzed data, and wrote the original manuscript.

## Figures and legend

**Fig. S1.**
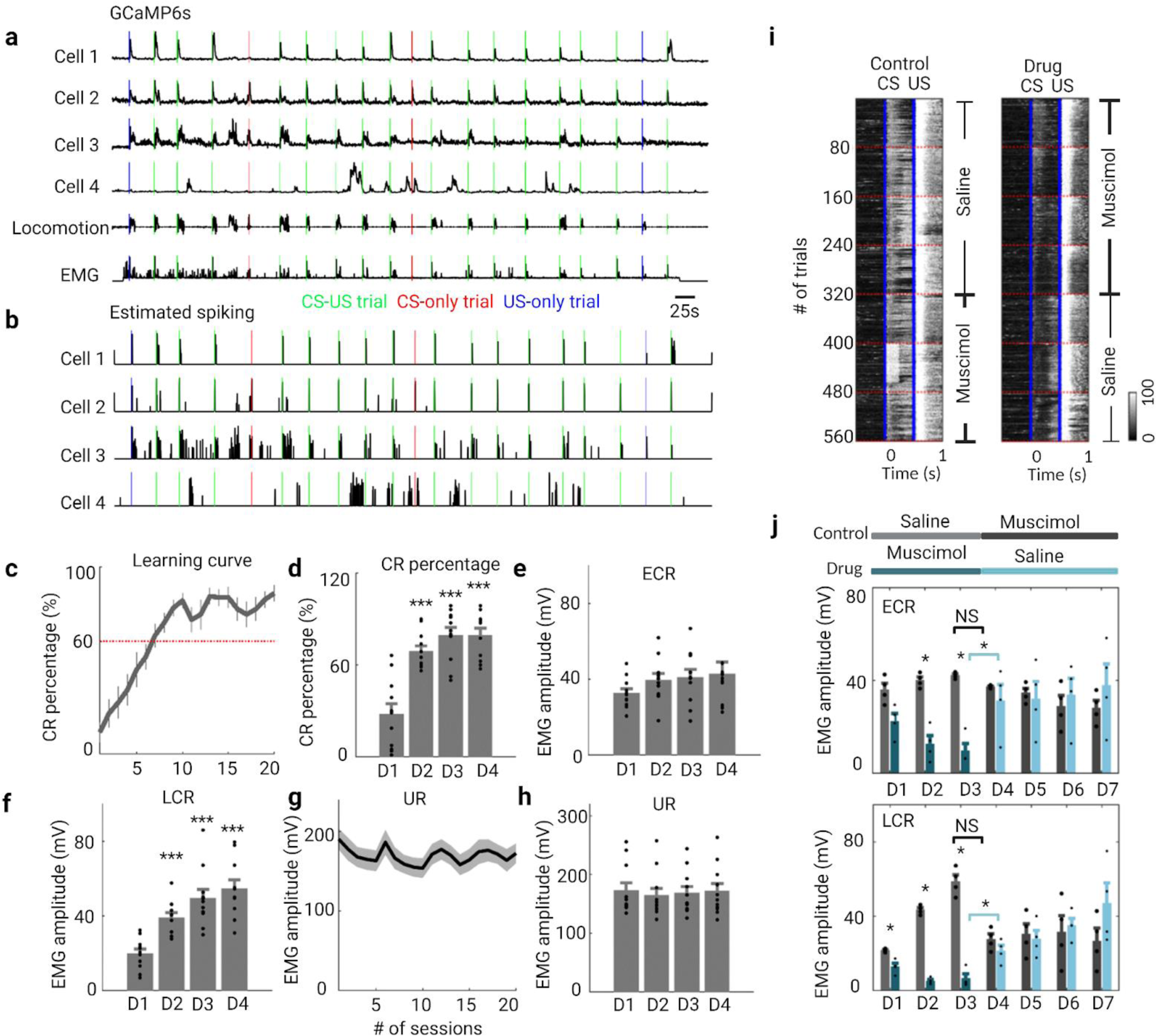
(Related to Fig. 1): Additional analyses of neural population responses and behavioral patterns. **a**, GCaMP6s signal (Δf/f) traces from four example L2/3 neurons, locomotion and EMG traces in the first session of TEC training. **b**, Estimated spiking of the example neurons in the first session of TEC training (green, CS-US; red, CS-only; blue, US-only). **c**, Evolution of conditional response percentage (CR%, top) and UR amplitude (bottom) across 20 sessions. Data are shown as mean ± s.e.m. Shaded areas represent s.e.m. **d∼h**, Behavior performance during training: CR percentage (n=11 mice; D1 vs. D2, ***P=9.1 × 10^−4^; D1 vs. D3, ***P=3.0 × 10^−4^; D1 vs. D4, ***P=3.0 × 10^−4^, Wilcoxon test), ECR (n=11 mice; D1 vs. D2, P=0.60; D1 vs. D3, P=0.84; D1 vs. D4, P=0.95, Wilcoxon test), LCR (n=11 mice; D1 vs. D2, ***P=5.0 × 10^−4^; D1 vs. D3, ***P=1.4 × 10^−4^; D1 vs. D4, ***P=1.1 × 10^−4^, Wilcoxon test) and UR (n=11 mice; D1 vs. D2, P=0.60; D1 vs. D3, P=0.84; D1 vs. D4, P=0.95, Wilcoxon test). Data are shown as mean ± s.e.m. Shaded areas represent s.e.m. **i**, Heat map of EMG responses from representative mice in control (left) and drug (right) groups during TEC learning (16 CS-US trials in one session, five sessions in a day shown in red dashed line; blue solid lines indicate onset of CS or US). The right side of each heat map indicates the treatment of muscimol or saline. **j**, Evolution of EMG amplitude (ECR, top; LCR, bottom) across 7 days (35 sessions) of training in the control group (saline was delivered during the first 3 days and then muscimol was delivered) and drug group (muscimol was delivered during the first 3 days and then saline was delivered) (n=4 in each group; ECR, control vs. drug in D1, P=0.057; control vs. drug in D2, *P=0.029; control vs. drug in D3, *P=0.029; control vs. drug in D4, P=0.89; control vs. drug in D5, P=0.89; control vs. drug in D6, P=0.49; control vs. drug in D7, P=0.49; control in D3 vs. control in D4, P=0.058; drug in D3 vs. drug in D4, P=0.029; LCR, control vs. drug in D1, *P=0.029; control vs. drug in D2, *P=0.029; control vs. drug in D3, *P=0.029; control vs. drug in D4, P=0.20; control vs. drug in D5, P=0.89; control vs. drug in D6, P=0.89; control vs. drug in D7, P=0.34; control in D3 vs. control in D4, P=0.057; drug in D3 vs. drug in D4, P=0.029; Wilcoxon test).

**Fig. S2.**
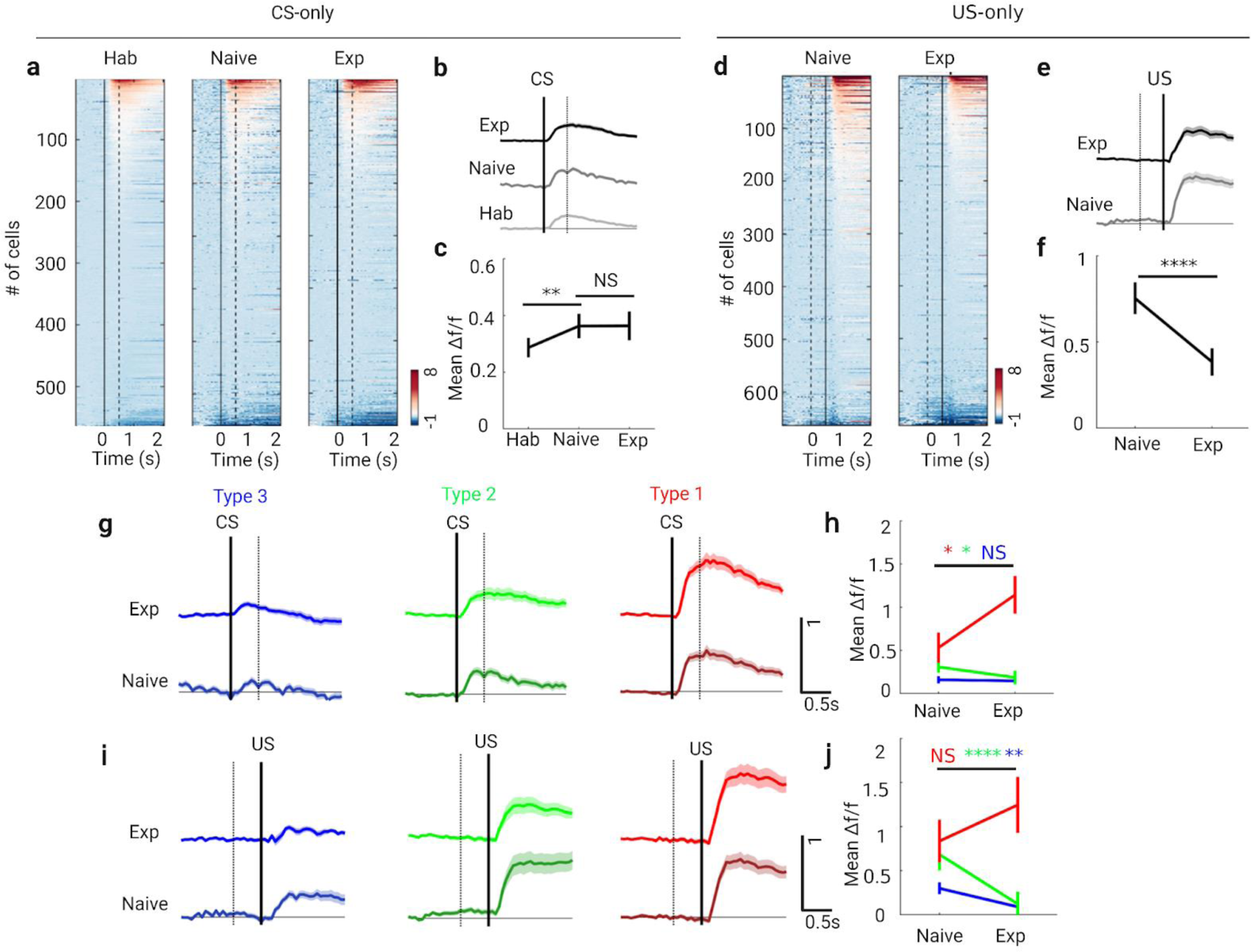
(Related to Fig. 1): Additional analyses of neural population responses and three types of PNs in CS-only and US-only trials. **a**, Heat maps of GCaMP6s signals (Δf/f) for PNs in CS-only trials at different stages of training (hab, n=566 neurons from 9 mice; Naive, n=663 neurons from 9 mice; Exp, n=665 neurons from 9 mice). Signals are aligned to the CS onset. **b**, Mean signal traces of PNs in CS-only trials at each learning stage. Data are shown as mean ± s.e.m. Shaded areas represent s.e.m. **c**, CR activity of PNs in CS-only trials at different training stages (hab vs. naive,**P=0.0061, naive vs. exp, NS P=0.99; Wilcoxon test). **d**, Heat maps of GCaMP6s signals (Δf/f) for PNs in US-only trials at different stages of training (Naive, n=663 neurons from 9 mice; Exp, n=665 neurons from 9 mice). Signals are aligned to the CS onset. **e**, Mean signal traces of PNs in US-only trials at each learning stage. Data are shown as mean ± s.e.m. Shaded areas represent s.e.m. **f**, UR activity of PNs in US-only trials at different training stages (naive vs. exp, ****P=3.62 × 10^−14^; Wilcoxon test). **g**, Mean signal traces of three subtypes of PNs in CS-only trials at each learning stage. Data are shown as mean ± s.e.m. Shaded areas represent s.e.m. **h**, CR activity of three subtypes of PNs in CS-only trials at different training stages (naive vs. exp in type 1,*P=0.011, naive vs. exp in type 2,*P=0.049, naive vs. exp in type 3, NS P=0.95; Wilcoxon test). **i**, Mean signal traces of three subtypes of PNs in US-only trials at each learning stage. Data are shown as mean ± s.e.m. Shaded areas represent s.e.m. **j**, UR activity of three subtypes of PNs in US-only trials at different training stages (naive vs. exp in type 1, NS P=0.84, naive vs. exp in type 2,**P=0.0014, naive vs. exp in type 3, ****P=3.04 × 10^−9^; Wilcoxon test).

**Fig. S3.**
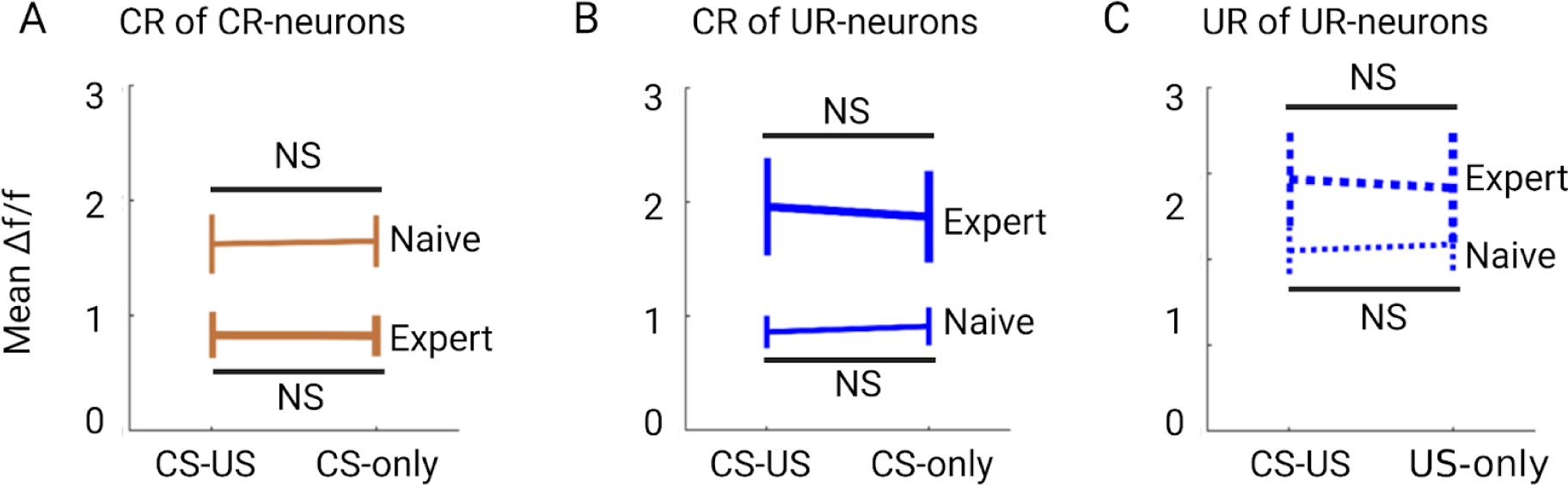
(Related to Fig. 2): Additional analyses of CR-neurons and UR-neurons in CS-only and US-only trials. **a**, Comparison of CR-neurons’ CS-induced responses (CR) in CS-US trials and CS-only trials at naive and expert stages (naive, NS P=0.88; expert, NS P=0.41; Wilcoxon test). **b**, Comparison of UR-neurons’ CS-induced responses (CR) in CS-US trials and CS-only trials at naive and expert stages (naive, NS P=0.36; expert, NS P=0.91; Wilcoxon test). **c**, Comparison of UR-neurons’ US-induced responses (UR) in CS-US trials and CS-only trials at naive and expert stages (naive, NS P=0.68; expert, NS P=0.18; Wilcoxon test). Data are shown as mean ± s.e.m.

**Fig. S4.**
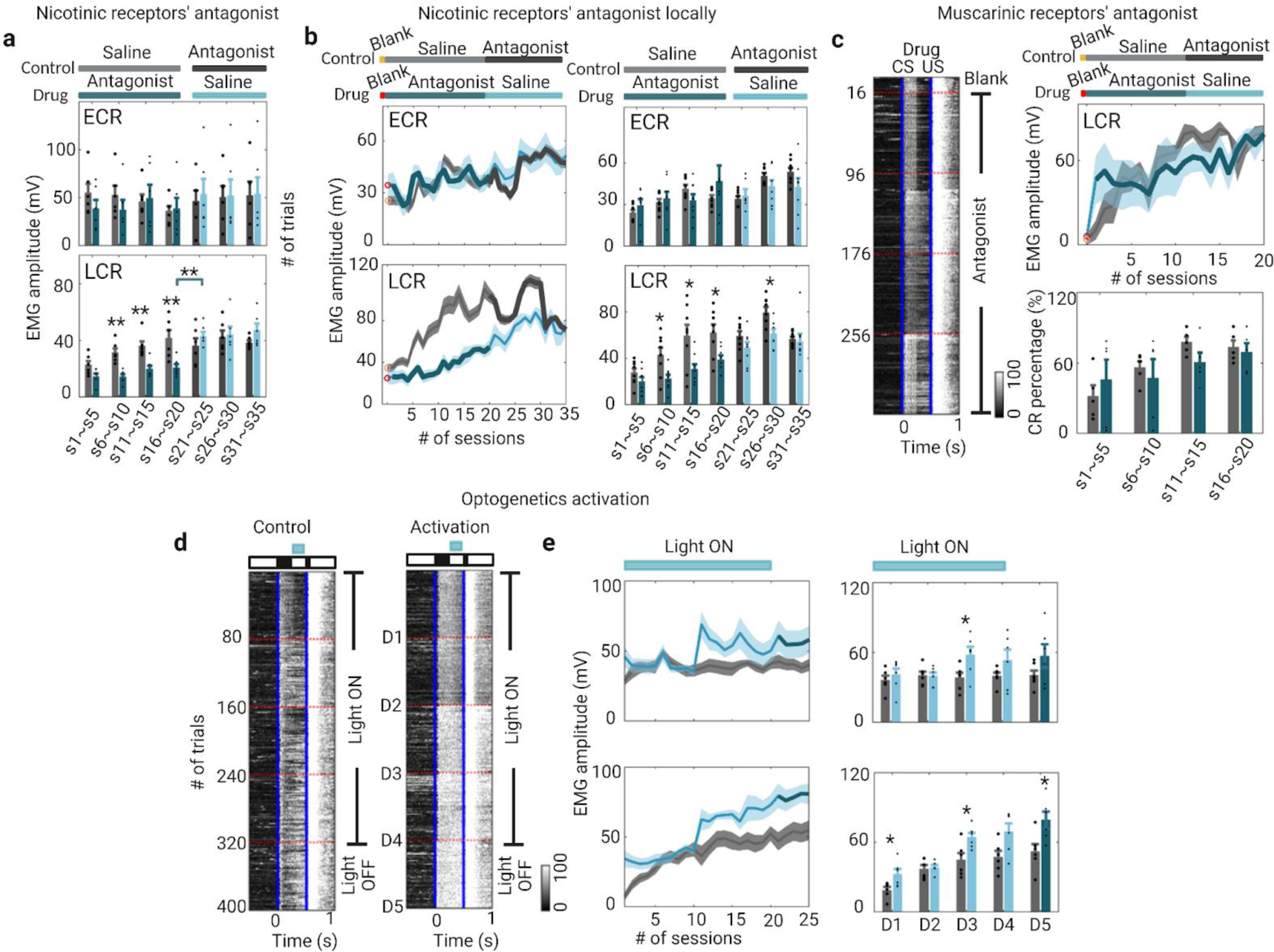
(Related to Fig. 3): EMG changes after pharmacological and optogenetic manipulation. **a**, EMG amplitudes (ECR, top; LCR, bottom) of nAChR antagonist and control groups (n=6 mice in each group; ECR, drug vs. control in s1∼s5, P=0.31, drug vs. control in s6∼s10, P=0.13, drug vs. control in s11∼s15, P=1, drug vs. control in s16∼s20, P=0.82, drug vs. control in s21∼s25, P=0.94, drug vs. control in s26∼s30, P=0.94, drug vs. control in s31∼s35, P=1; LCR, control in s1∼s5, P=0.093, drug vs. control in s6∼s10, **P=0.0022, drug vs. control in s11∼s15, **P=0.0087, drug vs. control in s16∼s20, **P=0.0087, drug vs. control in s21∼s25, P=0.31, drug vs. control in s26∼s30, P=0.82, drug vs. control in s31∼s35, P=0.13; Wilcoxon test). **b**, EMG changes after local treatment of nAChR antagonist and saline in S1. Left, evolution of EMG amplitude (ECR, top; LCR, bottom) across 35 sessions of training in the control group (One blank session in day 0, saline was delivered during the first 3 days and then nAChR antagonist was delivered) and drug group (One blank session in day 0, nAChR antagonist was delivered during the first 3 days and then saline was delivered) (n=8 in each group). Right, EMG amplitudes (ECR, top; LCR, bottom) of nAChRs antagonist and control groups (n=8 mice in each group; ECR, drug vs. control in s1∼s5, P=0.16, drug vs. control in s6∼s10, P=0.72, drug vs. control in s11∼s15, P=0.23, drug vs. control in s16∼s20, P=0.51, drug vs. control in s21∼s25, P=0.72, drug vs. control in s26∼s30, P=0.44, drug vs. control in s31∼s35, P=0.16; LCR, control in s1∼s5, P=0.19, drug vs. control in s6∼s10, *P=0.038, drug vs. control in s11∼s15, *P=0.049, drug vs. control in s16∼s20, *P=0.021, drug vs. control in s21∼s25, P=0.19, drug vs. control in s26∼s30, *P=0.038, drug vs. control in s31∼s35, P=0.23; Wilcoxon test).Data are shown as mean ± s.e.m. Shaded areas represent s.e.m. **c**, Left, heat map of EMG responses from muscarinic receptor (mAChR) antagonist treated mouse during TEC learning (16 CS-US trials in one session, five sessions in a day shown in red dashed line; blue solid lines indicate onset of CS or US). The right side of the heat map indicates the treatment of mAChR antagonist. Data are shown as mean ± s.e.m. Shaded areas represent s.e.m. Right, LCR (top) and CR percentage (bottom) changes of mAChRs antagonist and control group (n=5 mice in each group; ECR, drug vs. control in s1∼s5, P=0.69, drug vs. control in s6∼s10, P=0.69, drug vs. control in s11∼s15, P=0.10, drug vs. control in s16∼s20, P=0.75). **d**, EMG changes after optogenetic activation of cholinergic fibers in S1 (16 CS-US trials in one session, five sessions in a day shown in red dashed line; blue solid lines indicate onset of CS or US). The right side of the heat map indicates light ON and light OFF. **e**, Left, evolution of EMG amplitude (ECR, top; LCR, bottom) across 25 sessions of training in the control group (light ON during the first 3 days and then light OFF) and activation group (light ON during the first 3 days and then light OFF) (n=6 in each group). Right, EMG amplitudes (ECR, top; LCR, bottom) of activation and control groups (n=6 mice in each group; ECR, activation vs. control in D1, P=0.31, activation vs. control in D2, P=0.94, activation vs. control in D3, *P=0.041, activation vs. control in D4, P=0.31, activation vs. control in D5, P=0.39; LCR, activation vs. control in D1, *P=0.041, activation vs. control in D2, P=0.94, activation vs. control in D3, *P=0.015, activation vs. control in D4, P=0.065, activation vs. control in D5, *P=0.015; Wilcoxon test). Data are shown as mean ± s.e.m. Shaded areas represent s.e.m.

**Fig. S5.**
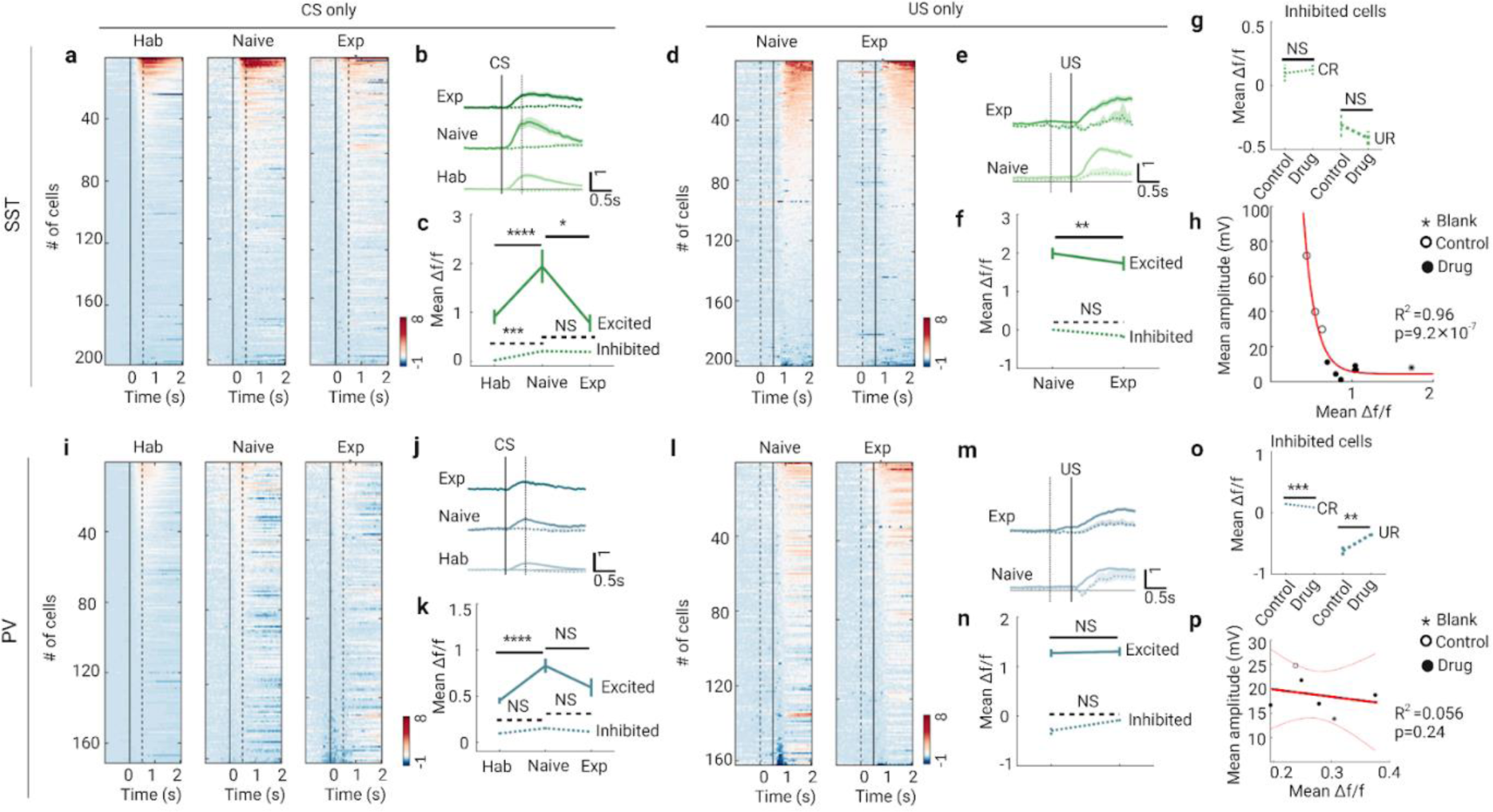
(Related to Fig. 4): Additional analyses of SST– and PV-INs. **a**, Heat maps of GCaMP6s signals (Δf/f) for SST-INs in CS-only trials at different stages of training (Hab, n=198 neurons from 7 mice; Naive, n=203 neurons from 7 mice; Exp, n=255 neurons from 7 mice). Signals are aligned to the CS onset. **b**, Mean signal traces of SST-INs in CS-only trials at each learning stage (solid line indicates the excited cells, dashed line indicates the inhibited cells). Data are shown as mean ± s.e.m. Shaded areas represent s.e.m. **c**, CR activity of excited (solid line) and inhibited (dashed line) SST-INs in CS-only trials at different training stages (CR of excited cells, hab vs. naive, ****P=1.97 × 10^−7^, naive vs. exp, *P=0.027; CR of inhibited cells, hab vs. naive, ***P=3.63 × 10^−4^, naive vs. exp, NS P=0.90; Wilcoxon test). **d**, Heat maps of GCaMP6s signals (Δf/f) for SST-INs in US-only trials at different stages of training (Naive, n=203 neurons from 7 mice; Exp, n=255 neurons from 7 mice). Signals are aligned to the CS onset. **e**, Mean signal traces of SST-INs in US-only trials at each learning stage (solid line indicates the excited cells, dashed line indicates the inhibited cells). Data are shown as mean ± s.e.m. Shaded areas represent s.e.m. **f**, UR activity of excited (solid line) and inhibited (dashed line) SST-INs in US-only trials at different training stages (UR of excited cells, **P=0.0050; UR of inhibited cells, NS P=0.053; Wilcoxon test). **g**, CR (fine line) and UR (bold line) activity of inhibited SST-INs in drug and control sessions (CR, NS P=0.44; UR, NS P=0.29; Wilcoxon test). **h**, Mean SST-INs’ signal versus mean EMG amplitude from CS onset to US onset in blank (asterisk), drug (solid circle) and control (hollow circle) sessions from a represented mouse (decay function fitting, n= 10 sessions from 1 mice). **i**, Heat maps of GCaMP6s signals (Δf/f) for PV-INs in CS-only trials at different stages of training (Hab, n=172 neurons from 5 mice; Naive, n=162 neurons from 5 mice; Exp, n=153 neurons from 5 mice). Signals are aligned to the CS onset. **j**, Mean signal traces of PV-INs in CS-only trials at each learning stage (solid line indicates the excited cells, dashed line indicates the inhibited cells). Data are shown as mean ± s.e.m. Shaded areas represent s.e.m. **k**, CR activity of excited (solid line) and inhibited (dashed line) PV-INs in CS-only trials at different training stages (CR of excited cells, hab vs. naive, ****P=6.72 × 10^−6^, naive vs. exp, P=0.099; CR of inhibited cells, hab vs. naive, NS P=0.11, naive vs. exp, NS P=0.67; Wilcoxon test). **l**, Heat maps of GCaMP6s signals (Δf/f) for PV-INs in US-only trials at different stages of training (Naive, n=162 neurons from 5 mice; Exp, n=153 neurons from 5 mice). Signals are aligned to the CS onset. **m**, Mean signal traces of PV-INs in US-only trials at each learning stage (solid line indicates the excited cells, dashed line indicates the inhibited cells). Data are shown as mean ± s.e.m. Shaded areas represent s.e.m. **n**, UR activity of excited (solid line) and inhibited (dashed line) PV-INs in US-only trials at different training stages (UR of excited cells, P=0.65; UR of inhibited cells, NS P=0.47; Wilcoxon test). **o**, CR (fine line) and UR (bold line) activity of inhibited PV-INs in drug and control sessions (CR, ***P=7.78 × 10^−4^; UR, **P=0.0033; Wilcoxon test). **p**, Mean PV-INs signal versus mean EMG amplitude from CS onset to US onset in blank (asterisk), drug (solid circle) and control (hollow circle) sessions from a represented mouse(Pearson correlation fit and 95% confidence bands, n= 6 sessions from 1 mice).

